# Using eDNA metabarcoding to establish targets for freshwater fish composition following river restoration

**DOI:** 10.1101/2022.05.26.493668

**Authors:** Gen Ito, Hiroshi Yamauchi, Miwa Shigeyoshi, Kousuke Ashino, Chie Yonashiro, Maki Asami, Yuko Goto, Jeffrey J. Duda, Hiroki Yamanaka

## Abstract

Establishing realistic targets for fish community composition is needed to assess the effectiveness of river restoration projects. We used environmental DNA (eDNA) metabarcoding with MiFish primers to obtain estimates of fish community composition across 17 sites upstream, downstream and within a restoration mitigation project area (Kaihotsu– Kasumi) located in the Shigenobu River system, Ehime Prefecture, Japan. We evaluate the benefits of using eDNA to quickly, sensitively, and extensively gather data to establish existing fish community composition in the restoration area, as well as potential future short-term, medium-term, and long-term targets of species assemblages that could realistically emerge following dispersal into the project area from upstream and downstream populations. We compare results from eDNA metabarcoding with species lists obtained from contemporaneous capture surveys and historical information. Nonmetric multidimensional scaling plots of community composition obtained from eDNA surveys showed that the Kaihotsu–Kasumi restoration area and surrounding river reaches were divided into three clusters: upper reaches, middle and lower reaches, and estuarine reaches. The Kaihotsu– Kasumi restoration area sites were included in the group containing the middle and lower reaches of the inflow and outflow rivers that were near the restoration area. We detected a total of twenty-six species in this group, twenty-one native species and five non-native species. Therefore, these native species were considered suitable as short-term target species with high potential for dispersal into Kaihotsu–Kasumi restoration area. By comparison, only 14 species would have been selected as target species based on capture surveys and historical literature. One factor increasing the resolution of our eDNA surveys was our ability to identify the presence of intraspecific lineages of *Misgurnus anguillicaudatus* (Clades A and B), which were missed by the capture surveys. These results indicate that the eDNA metabarcoding method can provide more comprehensive and realistic short-term target species estimates than capture surveys, as well as provide higher resolution monitoring through intraspecific lineage detection.

## Introduction

An imperative for aquatic biodiversity is to identify the organisms (hereafter “target species”) that can benefit from restoration actions (Morimoto and Kameyama, 2001; Wiens and Hobbs 2015). When restoring connectivity to river habitats, most projects rely upon aquatic organisms naturally dispersing into restored areas from connected waters. Setting targets of potential species assemblages following restoration actions is needed to assess project success (Beechie et al. 2008, Roni et al. 2008). Such understanding also provides information about the likelihood of project success, because the pool of target species and their metapopulation structure is a primary determinant of how fish species respond to ecosystem restoration (Huxel and Hastings 1999; Stoll et al. 2013; Stoll et al. 2014). However, it can be difficult to objectively identify which species from nearby contributing populations could be part of the reassembled target species pool. One approach relies on past to present spatial distribution data of organisms in neighboring areas based on capture surveys, but there are several issues that limit the utility of this approach. If historical spatial distribution data are used, the risk of including large areas where target species cannot disperse into exists, as well as the risk of including extinct or extirpated species. If recent spatial distribution data are used, it may not be of the proper scope (especially in terms of spatial coverage and resolution) to estimate the full list of potential target species. In addition, there may be no available spatial distribution data for nearby areas.

Several methods exist to estimate the presence of species in aquatic habitats using environmental DNA (eDNA). Such methods are desirable because of their sensitivity and ability to detect cryptic or rare organisms, often at a lower cost than traditional capture surveys which in turn allows for increased spatial and temporal sampling effort (Jerde et al., 2013; Oka et al., 2021; Sakata et al., 2017; Sigsgaard et al., 2015; Yamamoto et al., 2017). Many studies have shown the usefulness of eDNA methods for determining the spatial distribution and diversity of fish species (e.g., Doi et al., 2021; Nakagawa et al., 2018; Oka et al., 2021; Yamamoto et al., 2016, 2017) and that eDNA methods can be used to monitor rare species with low densities and to provide greater sensitivity in monitoring after removing artificial instream barriers (e.g., Duda et al., 2021; Muha et al., 2021; Sakata et al., 2017). In particular, the eDNA metabarcoding method can simultaneously detect many fish species at each location (e.g., Nakagawa et al., 2018; Yamamoto et al., 2017). Because the eDNA metabarcoding method can easily obtain spatial distribution data for current fish species, it can provide objective and realistic targets of target species that could assemble in restored habitats.

Freshwater fishes are a major target species for restoration mitigation in rivers, but the presence of cryptic non-native populations that cannot be distinguished by morphology is a problem for biodiversity conservation (Mabuchi et al., 2008; Watanabe et al., 2017). eDNA is a promising tool for detecting cryptic non-native populations (Uchii et al., 2016). MiFish primers, commonly used in eDNA metabarcoding studies (Miya et al., 2015), can detect fishes at the species level in most cases, but the resolution of intraspecific lineages is not well discussed. The ability to distinguish between native and non-native lineages using eDNA metabarcoding methods would be useful in target species selection for restoration and conservation programs. Three lineages of *Misgurnus anguillicaudatus* in the Japanese archipelago, which have been difficult to identify using morphological characteristics, are now identifiable using phylogenetic methods: two native lineages (Clades A and C sensu Koizumi et al., 2009) and a non-native lineage from China (Clade B sensu Koizumi et al., 2009). The native *M. anguillicaudatus* is categorized as *near threatened* by the Ministry of the Environment Government of Japan (2020). However, the non-native lineage has been found to be distributed throughout the Japanese archipelago (Morishiuma et al., 2008; Koizumi et al., 2009), which has a negative impact on the conservation of these native lineages. Native and non-native lineages of *M. anguillicaudatus* can be distinguished in the cytochrome *b* and control regions of mitochondrial DNA (Morishima et al., 2008; Koizumi et al., 2009). Therefore, MiFish for the 12S rRNA region of mitochondrial DNA may also distinguish them.

The Shigenobu River flows 36 km into the Seto Inland Sea, and wetlands with springs as their source of water exist along the banks of the middle reaches of the river. In the middle reaches of the Shigenobu River, including wetlands, good habitats (e.g., natural riverbeds and riverbanks) have decreased or deteriorated due to urbanization and levee construction in recent years (Ministry of Land, Infrastructure, Transport and Tourism. Shikoku Regional Development Bureau, 2008). Therefore, restoration mitigation was conducted by the Ministry of Land, Infrastructure, Transport and Tourism from 2014 to 2019 for Kaihotsu–Kasumi, one of the wetlands located in the middle reaches of the Shigenobu River, to restore the spring area (Water intake through a water pipeline from 500m upstream) and improve its connectivity with the mainstream (Removal of 0.5-1.0m levee and installation of pool-and-weir fishway). On the Shigenobu River, there were records of capture surveys conducted every five years from 1994 to 2021 (Ministry of Land, Infrastructure, Transport and Tourism, http://www.nilim.go.jp/lab/fbg/ksnkankyo/index.html, accepted 1 April 2022), as well as information on the distribution of endangered species (Ehime Prefecture, Japan, 2014). Given the regional species pool identified from prior capture surveys, a technical working group advised by experts and discussions with local residents created a target species list that could potentially assemble following restoration mitigation of Kaihotsu–Kasumi (hereafter “target species by the working group”, Table S1). The working group also identified potential response times as short-term (restoration of about 3 years), medium-term (restoration of about 20 years), and long-term (more than 20 years restoration) target species. In addition, monitoring was conducted in 2020 using both capture and eDNA metabarcoding surveys. The Kaihotsu–Kasumi restoration area of the Shigenobu River is a suitable location for testing whether the application of eDNA methods provides additional benefits selecting target species to track the effectiveness of restoration mitigation.

In this study, we examined the usefulness of eDNA metabarcoding to determine potential short-term target species assemblages using eDNA metabarcoding monitoring data collected from the Kaihotsu–Kasumi restoration area and neighboring habitats upstream and downstream. Additionally, we examined whether intraspecific lineages of an endangered native and non-native *M. anguillicaudatus* could be differentiated using DNA metabarcoding methods using MiFish primers.

## Methods

### Water sampling

Water samples were collected from Kaihotsu–Kasumi restoration area (5 sites: K1−5), the outflow river Shigenobu River (9 sites: S1−9), and the rivers (Uchi and Sagawa rivers) flowing into the Kaihotsu–Kasumi restoration area (3 sites: U1−3) from October 26 to October 27, 2020 (Fig. 1; Table S2). The basic water sampling protocol followed Minamoto et al. (2021), with some modifications. Single-use plastic cups were used to collect water samples, and 500 ml of surface water was collected per site. After water sampling, 500 μl of 10% benzalkonium chloride (BAC) was added immediately to inhibit DNA degradation (Yamanaka et al., 2017). Each water sample was filtered in the field by repeatedly loading a 50 ml water by syringe with a 0.45 μm pore size Sterivex filter. At the end of filtration, the Sterivex was plugged with sample water containing BAC to preserve eDNA.

**Fig. 1.**
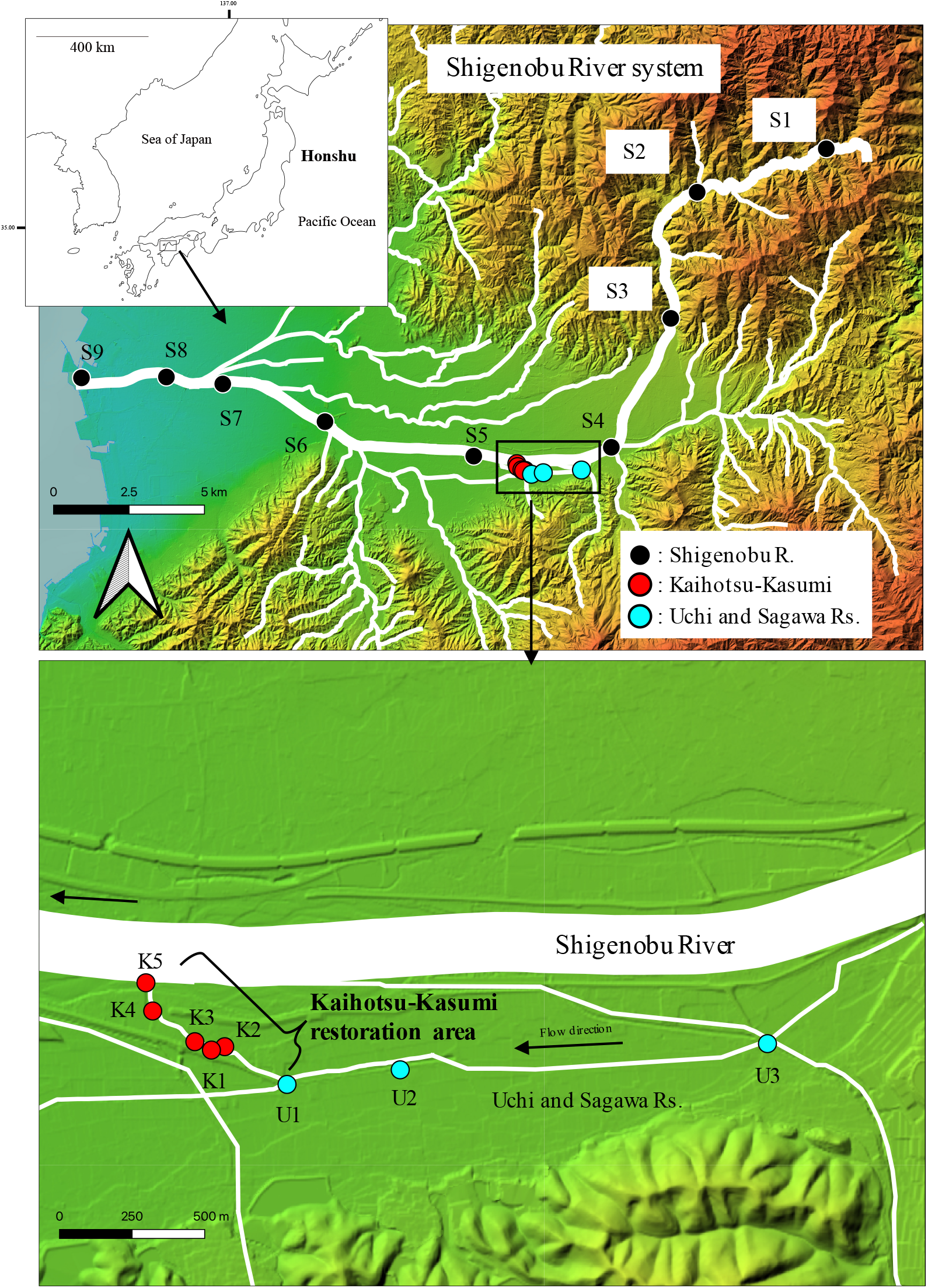
Water sampling sites in the Shigenobu River system. Each circle symbol indicates sampling sites where eDNA samples were collected. Concurrent fish capture surveys were also conducted at the 5 sites (K1-K5) of the Kaihotsu-Kasumi restoration area. This altitude map was used with permission from the Geospatial Information Authority of Japan (https://maps.gsi.go.jp/).

As a field negative control (FNC), 500 ml of ultrapure water was filtered using the same method as above after water sampling from three areas. After filtration, the samples were transported to the laboratory at room temperature, and the Sterivex filters were stored in a – 20 °C freezer on October 28, 2020.

### DNA extraction and amplification

DNA extraction was performed using the DNeasy Blood and Tissue Kit (Qiagen, Hilden, Germany), following Miya et al. (2016), with some modifications following Hirohara et al. (2021).

PCR was performed according to Miya et al. (2015), and the partial 12S rRNA region of mitochondrial DNA was amplified by the first PCR using the MiFish-6N-U-F/R primer pair. The 1st PCR was performed in a 12 μl reaction volume containing 0.24 μl of KOD Fx Neo (Toyobo, Osaka, Japan), 6 μl of 2 × Buffer for KOD Fx Neo, 2.4 μl of 2 mM dNTPs, 0.7 μl each of 5 μM MiFish-6N-U-F/R, and 2 μl of 10 × diluted template DNA. The thermal cycle profile was 94°C for 2 min; 40 cycles of 98°C for 10 s, 65°C for 30 s, and 68°C for 30 s; and 68°C for 5 min. Eight PCR replications were used for each sample and a PCR non-template control (PCR–NTC) using ultrapure water. The PCR amplicons were treated with ExoSAP-IT (ThermoFisher Scientific, Waltham, USA). The PCR amplicons from each of the eight replications were aggregated by 5 μl to a total of 40 μl, and 3.6 μl of ultrapure water and 0.4 μl of ExoSAP-IT were added per sample and treated at 37°C for 15 min and 80°C for 15 min. The second PCR was performed to add MiSeq adaptor sequences and eight bp index sequences (Hamady et al., 2008) to both amplicon ends. The total reaction volume of the 2nd PCR was 12 μl containing 0.24 μl of KOD Fx Neo, 6 μl of 2 × Buffer for KOD Fx Neo, 2.4 μl of 2 mM dNTP, 0.7 μl each of 5 μM MiFish-6N-U-F/R, 1 μl of ultrapure water, and 1 μl of the 1st PCR amplicons. The thermal cycle profile was 95°C for 3 min; 10 cycles of 98°C for 20 s, 65°C for 15 s, and 72°C for 15 s; and 72°C for 5 min. A 2% agarose gel electrophoresis was performed to confirm the amplification of the 2nd PCR amplicons.

### Amplicon library and Miseq sequencing

The samples were aggregated in volumes ranging from 2 or 3 μl per sample, depending on the results of the electrophoresis (Table S3). AMPure XP (Beckman Coulter, Indianapolis IN, USA) was added to 0.8 μl per 1 μl of each sample and treated according to the manual. DNA was eluted with 60 μl of TE buffer. Then, 25 μl of the treated DNA solution was added to a 2% E-Gel Size Select (Thermo Fisher Scientific, Waltham, MA, USA) to retrieve the target band (ca. 370 bp). The collected DNA concentration was measured using a Qubit dsDNA HS assay kit (Thermo Fisher Scientific, Waltham, MA, USA). Finally, a bioanalyzer (Agilent Technologies, Santa Clara, CA, USA) was used to confirm that PCR amplicons of the desired size were obtained.

The libraries were sequenced using an Illumina MiSeq with a reagent kit v2 nano for 2 × 150 bp PE (Illumina, San Diego, CA, USA). Raw reads were demultiplexed into each sample by a preinstalled program in MiSeq, and the outputted fastq files were processed by an in-house bioinformatic pipeline that uses USEARCH v11.0.667 (Edgar 2010) as its core program as follows. 1) Pair-end reads were merged using the *fastq_mergepairs* function. 2) Primer sequences were removed from both ends of the reads using *stripleft* and *stripright* functions. 3) Short reads less than 100 bp and those with a Q-Score less than 30 were removed using *fastq_minlen* and *fastq_truncqual* functions, respectively. 4) Identical reads were dereplicated using the *fastx_uniques* function and treated as operational taxonomic units (OTUs, i.e., zero-radius OTUs) with appending the sum of read numbers. 5) Denoising was performed using the *unoise3* function, and the OTUs with read counts size less than 4 were removed. Moreover, the OTUs that were judged as chimera were removed. 6) Taxonomic identity was assigned to the OTUs as follows: The sequences of each OTU were used as queries against a local sequence database at the threshold level of 0.95 using the *usearch_global* function. The database was a compilation of mitochondrial 12S sequences of fish downloaded from the National Center for Biotechnology Information website (https://www.ncbi.nlm.nih.gov/) on January 15, 2020. The hit sequences retrieved from the database for each OTU were converted into phylogenic trees and visually checked to assign a taxonomic identity for the focal OTU. The granularity of the taxonomic assignment to the OTUs was mainly at the species level, but some OTUs were identified at the genus level when a sequence cluster, including the query sequence and similar sequences at 0.95 threshold, included more than one species. Taxa that were unlikely to be distributed around the Shigenobu River (mainland of the Japanese archipelago) based on known geographic distribution information (Hosoya, 2019) were excluded (e.g., *Acanthopagrus sivicolus, Luciogobius_ryukyuensis*, and *Tridentiger kuroiwae*). Nucleotide sequence data reported are available in the DDBJ Sequenced Read Archive under the accession numbers DRR404794– DRR404814.

### Distinguishing between intraspecific lineages

For *M. anguillicaudatus*, we tried to distinguish between intraspecific lineages using the following methods. First, a 124 bp 12S rRNA (MiFish) tree was constructed using the sequence data output as *M. anguillicaudatus* OTU and *M. mizolepis* OTU. These OTUs were clustered as similar sequences at 0.95 threshold from the local sequence database of fish mitochondrial 12S sequences downloaded from NCBI. The names of these OTUs were named in correspondence with the names of the sequences in the most closely related databases. In addition, we used *Lefua echigonia*, a related species, as an outgroup (LC069488). Next, we constructed a 1015 bp cyt *b* tree, referring to Koizumi et al. (2009), in which molecular phylogenetic analyses of *M. anguillicaudatus* in Japan were conducted. For phylogenetic analysis using cyt *b*, nucleotide sequence data of *M. anguillicaudatus* and *M. dabryanus* registered in DDBJ, EMBL, and GenBank by Koizumi et al. (2009) were used: Clade A (native lineage, AB473261, AB473312), Clade B (non-native lineage, AB473360, AB473346), Clade C (native lineage, AB473381, AB473354, AB473382), and *M. dabryanus* (AB473408, AY625701). The complete mitochondrial genome sequences included in the MiFish phylogenetic tree (AP011291, AP017654, HM856629, KC509900, KC823274, MF579257, KF771003, KJ939363, KM186181, KM186183, MG938589, MG938590, NC027448) were also added to the cyt *b* phylogenetic tree. In addition, we used *L. echigonia* as an outgroup (LC079000, Ito et al., 2019). Multiple alignments of nucleotide sequences were performed using CLUSTAL W (Thompson et al., 1994). Phylogenetic analyses were conducted using the neighbor-joining (NJ) method. The NJ analyses were performed using MEGA X (Kumar et al., 2018), with the *p*-distance. The reliability of each internal branch was evaluated using a bootstrap test with 1000 replicates.

### Community analysis

The differences in community composition were visualized using nonmetric multidimensional scaling (nMDS). Dissimilarity matrices were calculated based on the Jaccard index using the vegdist function in the vegan package. A similarity profile analysis (simprof function) in the clustsig package was used to identify the number of significant cluster groups based on community similarity, and PERMANOVA was conducted using the adonis function in the vegan package to test whether the cluster groups obtained significantly differed from each other. All of the above statistics were conducted using R version 4.1.2 (R Core Team 2021).

### Capture survey

Capture surveys in Kaihotsu–Kasumi restoration area were conducted on August 27 and 28, 2020, and on October 26 and 27, 2020 at the same five sites as the eDNA sampling sites. The capture survey was conducted using hand nets, net pots, and bottle traps, with permission from Ehime Prefecture. The capture survey on October 26 was conducted after the eDNA survey.

## Results

### Miseq sequencing

A total of 1,435,754 reads were obtained by Miseq run from 17 sampling sites, 3 FNCs, and 1 PCR–NTC. Upon completion of the data processing pipeline, 314 OTUs were identified as fish species comprising 817,797 reads. Species assignment automatically conducted by Usearch was manually confirmed to check the independence of each OTU. If the representative sequence of the focal OTU could not be identified at the species level, the OTU was identified at the genus level. Five reads of *Opsariichthys platypus* were detected from an FNC of the Shigenobu River; therefore, the equivalent read number was subtracted from the *O. platypus* reads detected from the other sites in the Shigenobu River to account for possible cross-contamination (Yamamoto et al., 2017). No read was detected from the PCR–NTC.

### Fish species detected in the samples

We identified a total of 38 fish taxa from 8 orders and 16 families in our eDNA sampling. This list excludes 6 fish taxa likely present because of human consumption and runoff of eDNA into the sample location (i.e., Process Type I error; Darling and Mahon 2011): *Etrumeus teres, Trachurus japonicus, Pseudocaranx dentex, Seriola quinqueradiata, Pagrus major*, and *Pleurogrammus azonus*. However, peripheral freshwater fishes inhabiting estuarine brackish or ephemeral freshwater areas (*Plotosus japonicus, Mugil cephalus, Plectorhinchus cinctus, Acanthopagrus schlegelii, Omobranchus punctatus, Luciogobius pallidus, Luciogobius* spp., *Acanthogobius flavimanus, Tridentiger trigonocephalus, Favonigobius gymnauchen, Gymnogobius breunigii, Gymnogobius scrobiculatus, Takifugu* spp.) were included in the results. A total of 36 species were detected at the 9 Shigenobu River sites while 10 species were detected at the 3 Uchi and Sagawa River, and 19 species were identified from the five Kaihotsu–Kasumi restoration area sites (Table S4).

MiFish and cyt *b* trees are shown in Fig. 2. In the MiFish tree, the sequences of *M. anguillicaudatus* OTU and *M. mizolepis* OTU differed. The *M. anguillicaudatus* OTU included three sequences, OTU 57, 259, and 283, and 1 mitogenome sequence (AP011291).

**Fig. 2.**
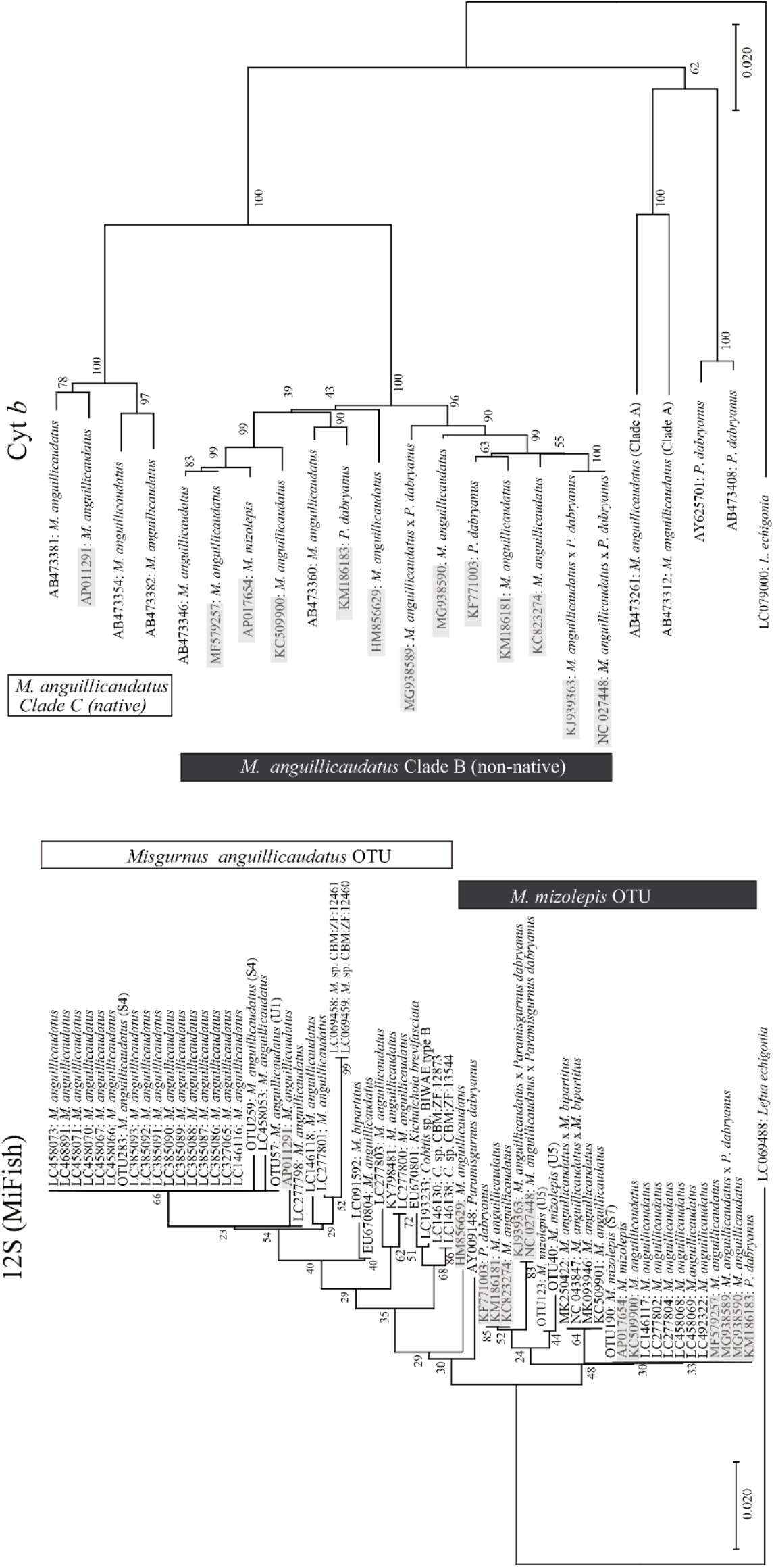
Neighbor-joining trees of 124 bp 12S rRNA (MiFish) gene (left) and 1015 bp cytochrome *b* gene (right) sequences of *M. anguillicaudatus*. The *numbers at the nodes* indicate bootstrap probabilities. The complete mitochondrial genome sequences are indicated by gray boxes. The *parentheses* of the MiFish tree indicate sampling sites.

The *M. mizolepis* OTU included three sequences, OTU 40, 123, and 190, and 12 mitogenome sequences. In the cyt *b* tree, three intraspecific lineages of *M. anguillicaudatus* (Clades A, B, and C), and *M. dabryanus* were distinguished with high bootstrap probability (100%), similar to Koizumi et al. (2009) and Perdices et al. (2016). Clade C (native lineage) contained one mitogenome sequence (AP011291), and Clade B (non-native lineage) contained the other 12 mitogenome sequences. No mitogenome sequences were identified in *M. dabryanus* or *M. anguillicaudatus* Clade A.

### Clustering based on community analysis

The community analysis results showed that the regional species pool detected across the 17 study sites was significantly differentiated into three cluster groups (Fig. 3; *P* < 0.001). Cluster 1 comprised three sites (S1−3) in the higher elevation portions (179−420 m) of the Shigenobu River. These sites had low species richness, with an average of 3.7 species detected per site (range 3−4) and a total of five taxa detected, including *Phoxinus oxycephalus jouyi, Oncorhynchus* spp, and *Rhinogobius flumineus*. Cluster 2 included five sites in the middle stream of the Shigenobu River (S4−8; elevation 7−100 m), and all three sites in the Uchi and Sagawa River (U1 − 3). Furthermore, all five sites in the Kaihotsu–Kasumi restoration area (K1−5) were also included in Cluster 2. Sites in cluster 2 had the highest average species richness of 14.5 (range 6−17) and a total of 26 taxa detected, including *Pseudorasbora parva, Gnathopogon elongatus, Oryzias latipes*, and *Odontobutis obscura*. Fourteen of these taxa were unique to Cluster 2. For Cluster 2, 23 taxa were identified from the Shigenobu River, 10 taxa from the Uchi and Sagawa River, and 19 taxa from Kaihotsu– Kasumi restoration area. Of these, *Tanakia limbata* and *Takifugu* spp. were detected only in Kaihotsu–Kasumi restoration area. Seven taxa, including *P. parva* and *Cobitis shikokuensis*, were detected only in the Shigenobu, Uchi, and Sagawa rivers (Table 1).

**Table 1.**
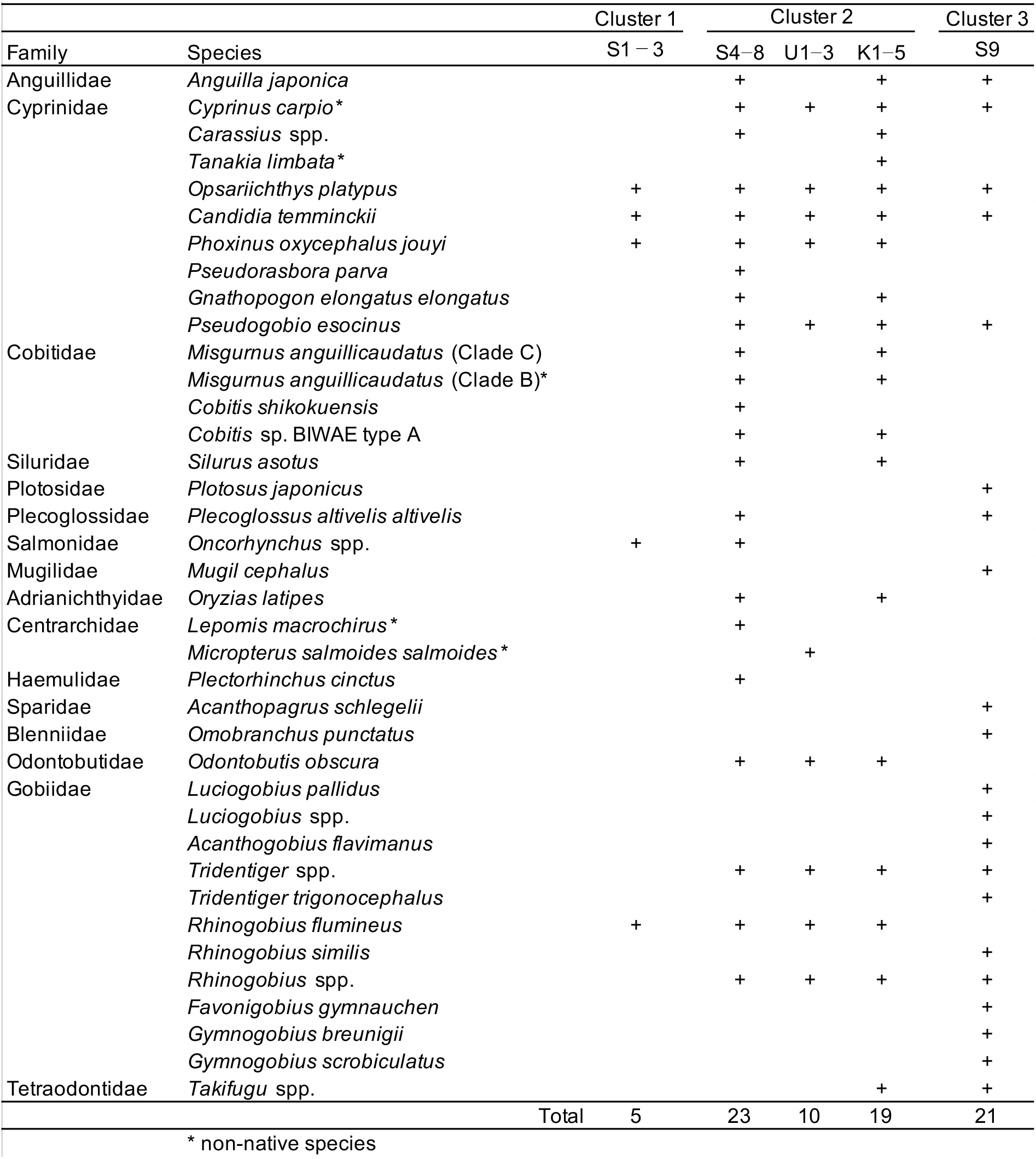
Fish species in each cluster based on nMDS. Cluster 1 comprised three sites (S1-3) in the higher elevation portions of the Shigenobu River. Cluster 2 included five sites in the middle stream of the Shigenobu River (S4-9), all five sited in the Kaihotsu-Kasumi restoration area (K1-5), and all three sites in the Uchi and Sagawa Rivers (U1-3). Cluster 3 comprised only one site (S9) in the estuary of the Shigenobu River.

**Fig. 3.**
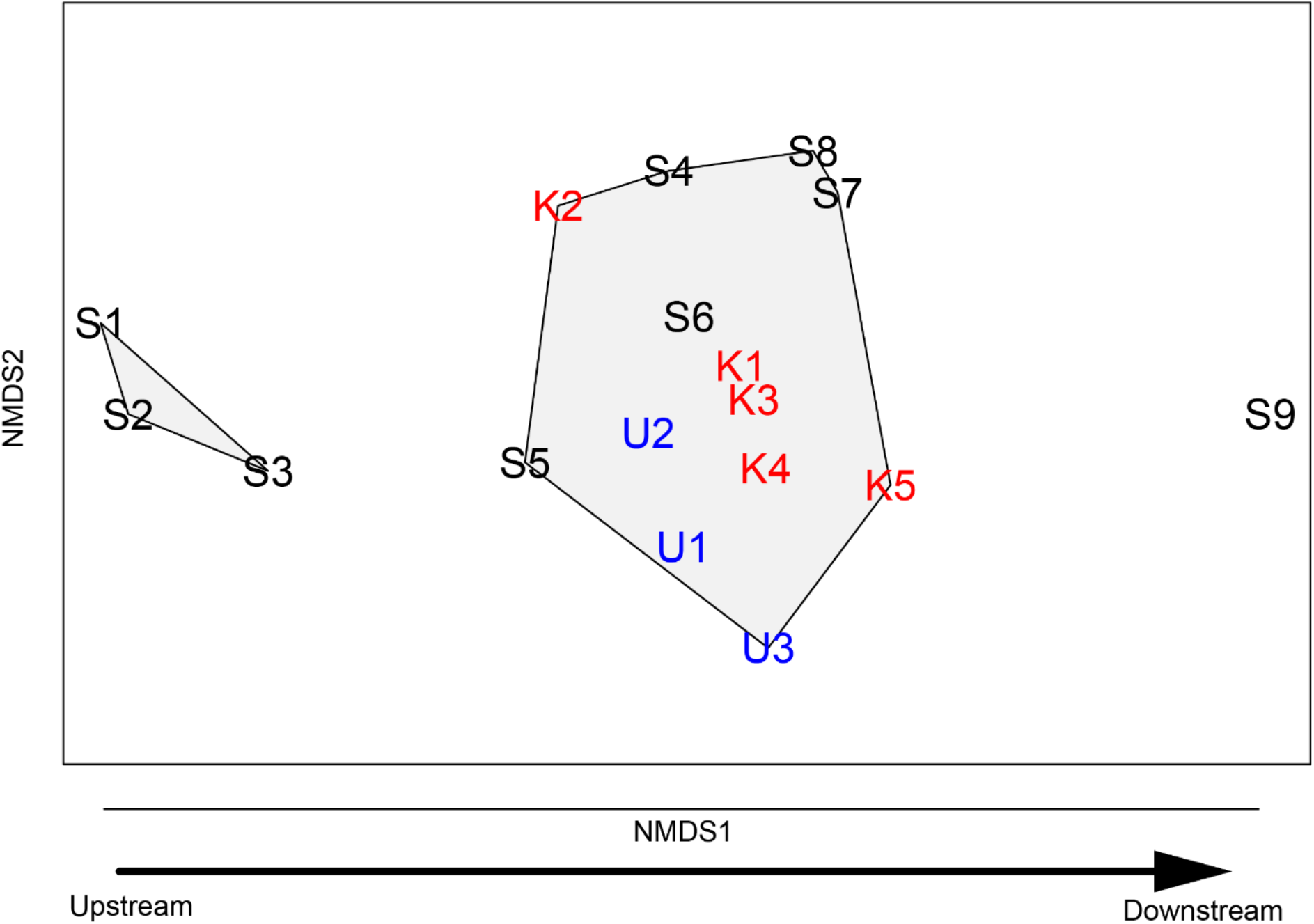
NMDS plot of fish species community compositions at different eDNA sampling sites. The community was significantly differentiated into three groups (PERMANOVA: *P* < 0.001). Site codes and text color follows map depicting sampling sites in the Shigenobu River (black), the Kaihotsu-Kasumi restoration area (red) and the Uchi and Sagawa Rivers (blue).

Cluster 3 comprised only one site (S9) in the estuary of the Shigenobu River (elevation −0.8 m), where 21 taxa were detected, including *Plotosus japonicus, Mugil cephalus*, and *Acanthopagrus schlegelii*. Twelve of these taxa were detected only in Cluster 3 (Table 1).

### Fish species detected by capture surveys

The capture surveys in 2020 identified 15 taxa in Kaihotsu–Kasumi restoration area (Table 2). *C. shikokuensis, Plecoglossus altivelis altivelis*, and *Lepomis macrochirus* were identified only in the capture surveys. Capture surveys failed to detect 5 taxa that were identified in eDNA sampling: *Anguilla japonica, Tanakia limbata, Gnathopogon elongatus elongatus, Silurus asotus*, and *Takifugu* spp. (Table 2). For *M. anguillicaudatus*, we were able to distinguish them as Clades B and C by eDNA surveys but were unable to distinguish both clades by capture surveys. *Rhinogobius nagoyae* and *Rhinogobius* sp. OR could be distinguished by capture surveys, but not by eDNA surveys. Fish species that could not be distinguished in one survey are shown in gray boxes in Table 2.

**Table 2.** Fish species confirmed by eDNA and capture surveys in Kaihotsu–Kasumi restoration area. Gray boxes are showed fish species that could not be identified using one of the techniques.

| Family | Species | eDNA | Capture |
| --- | --- | --- | --- |
| Anguillidae | *Anguilla japonica* | + |  |
| Cyprinidae | *Cyprinus carpio* * | + | + |
|  | *Carassius* spp. | + | + |
|  | *Tanakia limbata* * | + |  |
|  | *Opsariichthys platypus* | + | + |
|  | *Candidia temminckii* | + | + |
|  | *Phoxinus oxycephalus jouyi* | + | + |
|  | *Gnathopogon elongatus elongatus* | + |  |
|  | *Pseudogobio esocinus* | + | + |
| Cobitidae | *Misgurnus anguillicaudatus* (Clade C) | + | + |
|  | *Misgurnus anguillicaudatus* (Clade B)* | + |  |
|  | *Cobitis shikokuensis* |  | + |
|  | *Cobitis* sp. BIWAE type A | + | + |
| Siluridae | *Silurus asotus* | + |  |
| Plecoglossidae | *Plecoglossus altivelis altivelis* |  | + |
| Adrianichthyidae | *Oryzias latipes* | + | + |
| Centrarchidae | *Lepomis macrochirus* * |  | + |
| Odontobutidae | *Odontobutis obscura* | + | + |
| Gobiidae | *Tridentiger* spp. | + | + |
|  | *Rhinogobius flumineus* | + | + |
|  | *Rhinogobius nagoyae* | + | + |
|  | *Rhinogobius* sp. OR |  | + |
| Tetraodontidae | *Takifugu* spp. | + |  |
| Total |  | 19 | 17 |
| * non-native species |  |  |  |

## Discussion

We conducted eDNA sampling and metabarcoding analysis to assess the community composition of restored wetland areas and predict the target community composition based on beta diversity of fish throughout the adjacent riverscape. When compared with historical and contemporary field-based surveys of fish assemblages, results from metabarcoding analyses were similar once known false positives (Process type I error from species purchased commercially and brought into the surrounding communities for human food consumption) were removed from the dataset. Additionally, for some taxa that were challenging to identify to the species level in the field due using morphological characteristics, the genetic approaches allowed a greater taxonomic resolution to identify the pool of fish species present in the study area. Given the lower cost, similarity of target species lists compared with traditional survey methods, and higher taxonomic resolution for cryptic taxa, our results support the idea that eDNA techniques can be used to track fish communities following habitat restoration actions (Duda et al. 2020; Muha et al. 2021) and other similar environmental applications (e.g., DiBattista et al. 2022; Gold et al. 2021; Li et al. 2022).

All of the 38 taxa in 16 families of fishes detected by eDNA in this study were recorded in previous capture surveys in the Shigenobu River (Ministry of Land, Infrastructure, Transport and Tourism, http://www.nilim.go.jp/lab/fbg/ksnkankyo/index.html, accepted 1 April 2022). Therefore, the eDNA survey results were determined to be reasonable. The six taxa which could not be identified to the species level by eDNA was due to low interspecific variation in the MiFish region or introgression of mtDNA (e.g., *Carassius* spp.: Yamamoto et al., 2010; *Rhinogobius* spp.: Yamasaki et al., 2015). Therefore, we identified the following taxa by comparison with previous capture surveys (Ministry of Land, Infrastructure, Transport and Tourism, http://www.nilim.go.jp/lab/fbg/ksnkankyo/index.html, accepted 1 April 2022): *Carassius* spp. as *Carassius* sp. (Silver crucian carp) or/and *Carassius buergeri* (Giant golden crucian carp), *Oncorhynchus* spp. as *Oncorhynchus masou ishikawae* or/and *Oncorhynchus mykiss, Luciogobius* spp. as *Luciogobius guttatus* or/and *Luciogobius martellii, Tridentiger* spp. as *Tridentiger obscurus* or/and *Tridentiger brevispinis, Rhinogobius* spp. as *Rhinogobius nagoyae, Rhinogobius fluviatilis, Rhinogobius* sp. OR or/and *Rhinogobius tyoni*, and *Takifugu* spp. as *Takifugu niphobles* or/and *Takifugu poecilonotus*.

In this paper, to attempt to resolve intraspecific lineages using DNA metabarcoding methods using MiFish primers, we attempted to identify intraspecific lineages in *M. anguillicaudatus*. Based on the results of the Cyt *b* phylogenetic tree, including the mitogenome sequence, *M. anguillicaudatus* OTU is the native lineage (Clade C) and *M. mizolepis* OTU is the non-native lineage (Clade B). *M. anguillicaudatus* Clade B has also been recorded from the Shigenobu River system as a non-native lineage by capture surveys (Shimizu and Takagi, 2010). No matching sequences were found between *M. anguillicaudatus* and *M. mizolepis* OTUs; thus, both OTUs are genetically distinct. However, *M. dabryanus* (including *Paramisgurnus dabryanus* and *M. mizolepis*) and *M. anguillicaudatus* sequences were mixed in the *M. mizolepis* OTU. Molecular phylogenetic studies of mtDNA indicate that the *M. dabryanus* is not contained within *M. anguillicaudatus* Clades A and B, and shows a phylogenetic position closely related to *M. anguillicaudatus* Clade A and *M. nikolskyi* (Koizumi et al., 2009; Perdices et al., 2016). *M. anguillicaudatus* and *M. dabryanus* can be difficult to identify in the field using morphological characteristics. Therefore, the cause of the confusion of the sequence labeled “*M. dabryanus”* in *M. anguillicaudatus* Clade B could be the misidentification of *M. anguillicaudatus* as *M. dabryanus*. The sequence may also be a hybrid individual, since *M. anguillicaudatus* and *M. dabryanus* can hybridize (Okada et al., 2020). Intentional exclusion of sequences derived from misidentification and hybridization is difficult. Therefore, a phylogenetic tree that includes mitogenomic sequences would help identify intraspecific lineages.

Native lineage (Clade C) and non-native lineage (Clade B) of *M. anguillicaudatus* were identified from Kaihotsu–Kasumi restoration area using eDNA metabarcoding methods. The successful identification of native and non-native lineages of *M. anguillicaudatus* using eDNA is one of the advantages of introducing eDNA in monitoring. Future studies on the feasibility of identifying intraspecific lineages for a larger number of fishes will contribute to improving the resolution of monitoring using the eDNA metabarcoding method.

Multivariate analyses showed that the fish community structure of the Shigenobu River was significantly divided into three clusters (Clusters 1–3). Each cluster has an upstream (Cluster 1), middle and downstream (Cluster 2), and estuarine (Cluster 3) fish species assemblage, suggesting that the fish community assembles across a gradient of river environmental conditions (e.g., salinity, elevation, and gradient). The communities in Kaihotsu–Kasumi restoration area (K1–5) were included in Cluster 2, which showed a community structure associated with other mainstem sites located in the middle and lower reaches of the river. Thus, the fish community at Kaihotsu–Kasumi restoration area more closely resembles the middle and lower reaches of the Shigenobu River, which are geographically close to each other. Of the 26 taxa of fish detected at the Cluster 2 sites, 17 were common between Kaihotsu–Kasumi restoration area and neighboring waters (S4–8 and U1–3). The similarity between the fish communities in Kaihotsu–Kasumi restoration area and neighboring waters after connectivity was restored suggests that the natural dispersal of fishes inhabiting Kaihotsu–Kasumi restoration area from neighboring waters led to homogenization between these reconnected habitats (Rahal 2007). Kaihotsu–Kasumi restoration area is considered to provide a good fish habitat due to its high connectivity with neighboring waters.

Nineteen taxa were detected by eDNA in Kaihotsu–Kasumi restoration area, five of which were not detected in the capture surveys conducted during the same period. Fish surveys using eDNA tend to detect more species than capture surveys (e.g., Oka et al., 2021), and this study showed similar results. *Plecoglossus altivelis altivelis, C. shikokuensis*, and *L. macrochirus* were detected only in the capture surveys. Although optimization of eDNA metabarcoding methods is necessary to maximize detection of fish communities, a combination of capture surveys with eDNA approaches may be effective for a more accurate understanding of the fish community.

The fish community structure of Kaihotsu–Kasumi restoration area sample sites resembled the middle and lower reaches of the river (Fig. 3). Therefore, 21 taxa could be selected as short-term target species for Kaihotsu–Kasumi restoration area, excluding non-native species (*Cyprinus carpio, Tanakia limbata, M. anguillicaudatus* non-native lineage, *L. macrochirus, Micropterus salmoides*; Hosoya, 2019) out of the 26 taxa included in Cluster 2. The target species not identified by eDNA survey in Kaihotsu–Kasumi restoration area were *Pseudorasbora parva, Oncorhynchus* spp., and *Plectorhinchus cinctus. P. parva* is categorized as Near Threatened by Ehime Prefecture, Japan (2014). The slow flow of Kaihotsu–Kasumi restoration area is considered a suitable habitat and breeding environment for *P. parva*. In the future, efforts should be made to facilitate the natural dispersal of *P. parva* into the restored Kaihotsu–Kasumi restoration area reach. *Oncorhynchus* spp. and *P. cinctus* are found in the upper reaches of rivers and estuaries, respectively (Hosoya, 2019), therefore Kaihotsu–Kasumi restoration area located in the middle reaches of rivers may not be suitable habitat for both species. These two species may not be appropriate as short-term target species for Kaihotsu–Kasumi restoration area, further studies of selection methods, including fish preference by habitat, will bring us closer to a more realistic selection of short-term target species.

In this paper, we identified list of short-term target species expected to reoccupy the Kaihotsu-Kasumi reach after restoration mitigation using the eDNA metabarcoding method. As a result, 21 short-term target species were selected. A similar exercise of developing a target species list was conducted by a working group using traditional capture-based survey methods, with 9, 5, and 2 species identified as short-term (within 3 years), medium-term (within 20 years), and long-term (>20 years), respectively (Table S1). Of these, 14 of the short- and medium-term target species were among the short-term target species detected by eDNA (this study). Seven new species were included in the short-term target species established by eDNA. Therefore, the decision of short-term target species using the eDNA metabarcoding method is more effective than capture surveys in selecting fish species that are expected to recover in the short term in a more comprehensive manner, probably because eDNA is more sensitive to the current fish community than capture surveys.

## Credit author statement

Gen Ito: Data analysis, Writing – original draft, Writing – review and editing. Hiroshi Yamauchi: Planning and conducting field surveys, Writing – review and editing. Miwa Shigeyoshi, Kousuke Ashino and Chie Yonashiro: Planning and conducting field surveys. Maki Asami and Yuko Goto: laboratory processing. Jeff Duda: Writing– review and editing. Hiroki Yamanaka: Planning and conducting field surveys, Data analysis, Writing – review and editing, Conceptualization, Funding acquisition.

## Acknowledgements

We express sincere thanks to members of the Civil Engineering Division1, Matsuyama River National Highway Office, Shikoku regional development bureau for their help with the field surveys. Any use of trade, firm, or product names is for descriptive purposes only and does not imply endorsement by the U.S. Government.

